# Identifying Model-Based and Model-Free Patterns in Behavior on Multi-Step Tasks

**DOI:** 10.1101/096339

**Authors:** Kevin J. Miller, Carlos D. Brody, Matthew M. Botvinick

## Abstract

Recent years have seen a surge of research into the neuroscience of planning. Much of this work has taken advantage of a two-step sequential decision task developed by Daw et al. (2011), which gives the ability to diagnose whether or not subjects’ behavior is the result of planning. Here, we present simulations which suggest that the techniques most commonly used to analyze data from this task may be confounded in important ways. We introduce a new analysis technique, which suffers from fewer of these issues. This technique also presents a richer view of behavior, making it useful for characterizing patterns in behavior in a theory-neutral manner. This allows it to provide an important check on the assumptions of more theory-driven analysis such as agent-based model-fitting.

## Introduction

It has long been known that humans and animals construct internal models of the dynamics and contingencies in their environment (“cognitive maps”; Tolman 1948), and use these models to inform their decisions. Such “model-based” decision-making behavior is defined in learning theory (Sutton and Barto 1998) as planning, and is distinguished from simpler “model-free” strategies. Despite great progress in recent years elucidating the neural mechanisms of simple decision-making, the neuroscience of planning is still in its infancy. One important reason for this is that traditional planning tasks, such as outcome devaluation (Adams and Dickinson 1981), are limited to producing only one or a few instances of planned behavior per subject, sharply limiting the types of experimental designs that can be brought to bear. Recently, Daw et al., (2011) have developed a two-step sequential decision-making task that lifts this limitation, obtaining many hundreds of decisions per session, along with a behavioral readout of whether or not those decisions were planned.

The development of this task constituted a major step forward in the neuroscience of model-based behavior, rendering planning accessible for the first time to a wide range of new experimental designs. Recent work has used it to characterize the neural correlates of planning (Daw et al. 2011; Doll et al. 2015; Deserno et al. 2015), the structures and neurotransmitters involved (Wunderlich, Smittenaar, and Dolan 2012; Smittenaar et al. 2013; Worbe et al. 2015), factors that promote or interfere with planning (Otto, Raio, et al. 2013; Sebold et al. 2014; Eppinger et al. 2013; Otto, Gershman, et al. 2013; Radenbach et al. 2015; Voon et al. 2015), and relationships between planning and other cognitive phenomena (Skatova, Chan, and Daw 2013; Gillan et al. 2015; Schad et al. 2014; Sebold et al. 2016; Otto et al. 2015). In addition, variants of the task for animal subjects are being developed (Miranda, Malalasekera, Dayan, & Kennerly, 2013, *Society for Neuroscience Abstracts*; Akam, Dayan, & Costa, 2013, *Cosyne Abstracts*; Groman, Chen, Smith, Lee, & Taylor, 2014, *Society for Neuroscience Abstracts*; Hasz & Redish, 2016, *Society for Neuroscience Abstracts*; Miller, Botvinick, & Brody, in prep), opening a wide range of new possibilities for experiments investigating the neural mechanisms of planning.

Crucial to much of this is a set of analysis tools used to quantify the extent to which a behavioral dataset shows evidence of planning vs. the extent to which it shows evidence of having been produced by a model-free algorithm. Two such tools are commonly in use, both originating with the seminal work on this task (Daw et al., 2011). The first of these tools is a simple analysis of one-trial-back stay/switch choice behavior which is relatively theory neutral. Particular patterns are expected to appear in this analysis which are characteristic of planning vs. model-free systems. The second tool involves fitting the data with a theoretically motivated agent-based computational model, and interpreting the best fit parameters which result. Given the expanding popularity of the two-step task, it is important to thoroughly vet these tools, and determine the extent to which they provide a complete and accurate picture of the generative process which gave rise to a behavioral dataset, including in modified versions of the task optimized for animal subjects.

Here, we generate synthetic datasets for which the “ground truth” generative process is known, and subject them to analysis using both of these tools. We show that the one-trial-back analysis is subject to important confounds, rendering it poorly suited to the task of quantifying model-based vs. model-free patterns in behavior in several cases. We show that agent-based model-fitting analyses perform as expected in cases where their assumptions are met, but can fail dramatically when they are not. We introduce a novel theory-neutral tool for characterizing behavioral data on this task (the “many-trials-back” analysis). This tool used on its own is resistant to some (but not all) of the limitations associated with the one-trial-back analysis. It also provides a rich description of the qualitative patterns that are present in behavior, making it an effective tool for checking the assumptions of theory-driven approaches such as explicit model fitting.

We caution against the use of the one-trial-back analysis as a means of quantifying behavioral patterns, as well as the application of model-fitting analyses to novel datasets in the absence of model comparison or validation by comparison to theory-neutral methods. We introduce the many-trials-back method both as an independent way of quantifying patterns in behavior, and recommend an approach to data analysis that combines this tool with explicit computational modeling.

## Methods

### Two-Step Behavioral Task

We generated synthetic behavioral data from the two-step Markov decision task developed by Daw et al. (2011), the structure of which is outlined in Figure 1. In the first step of this task, the agent must select between two actions (A&B in figure 1). In the second stage, the agent is presented with the choice between one of two pairs of actions (C&D or E&F in figure 1). The selection of a second-stage action results in either a reward or an omission.

**Figure 1:**
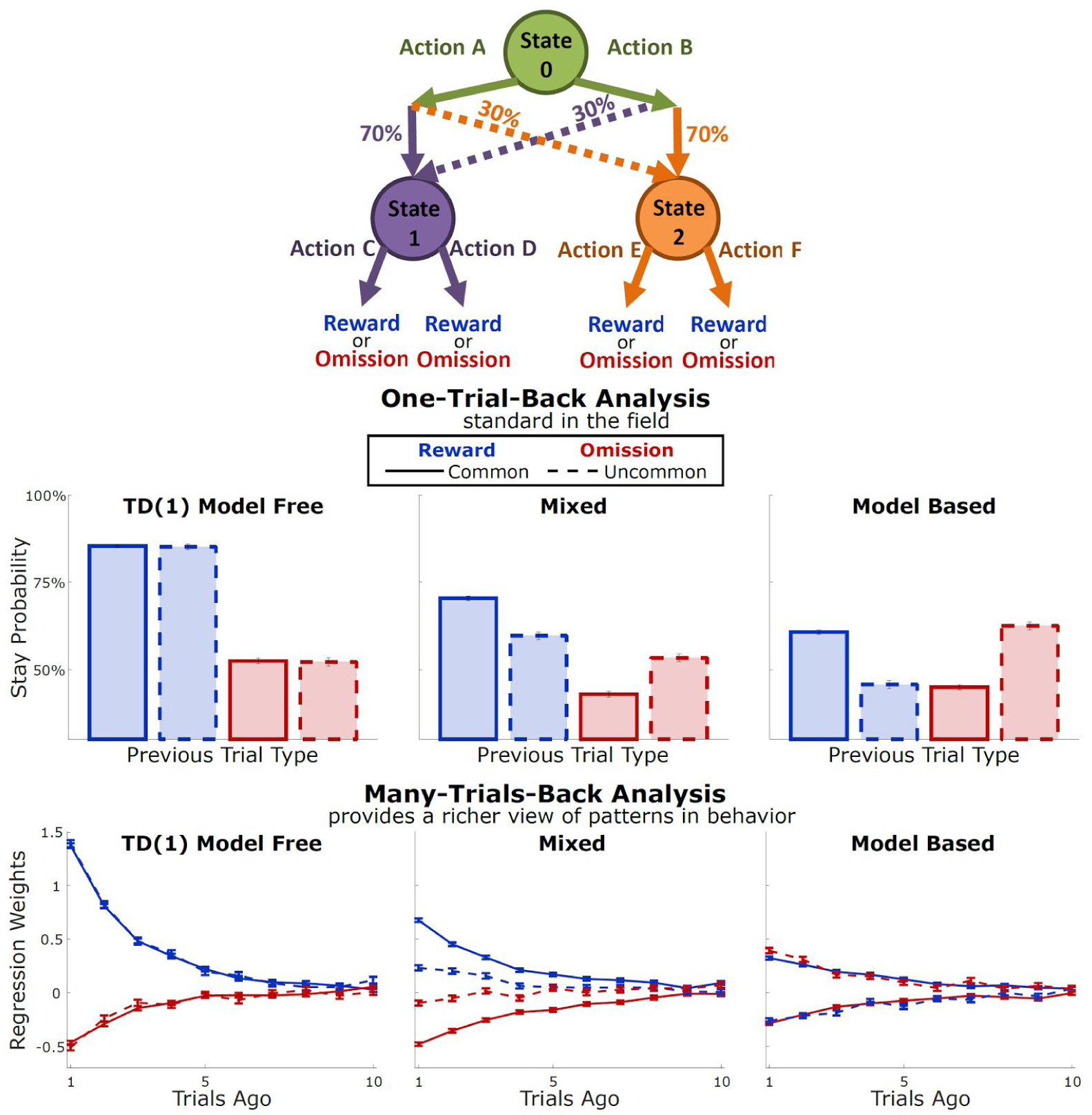
Model-free and model-based patterns of behavior on the Daw et al., (2011) version of the two-step task.

Which pair of second-stage choices was available on a particular trial depended probabilistically on the action selected in the first stage. Selection of Action A led to C&D becoming available 70% of the time and E&F becoming available 30% of the time, while selection of Action B led to C&D 30% of the time and E&F 70% of the time. The probability of a reward for each second-stage action was initialized randomly between 0 and 1, and evolved after each trial according to a gaussian random walk (μ = 0, σ = 0.025, as in Daw et al. 2011), with bounds at 0.25 and 0.75.

We also consider a simplified version of the two-step task, optimized for use with animal subjects, which is outlined in figure 2. This task eliminates the second choice. Selection of Action A at the first step led to Action C (only) becoming available 80% of the time, and Action D 20% of the time, while selection of Action B led to C or D with opposite probabilities. At all times, the probability of a reward following one second-step action was 80%, and following the other 20%. After each trial, there was a 2% chance that the reward probabilities would reverse, with the constraint that they could not reverse if they had reversed previously within the last 10 trials.

**Figure 2:**
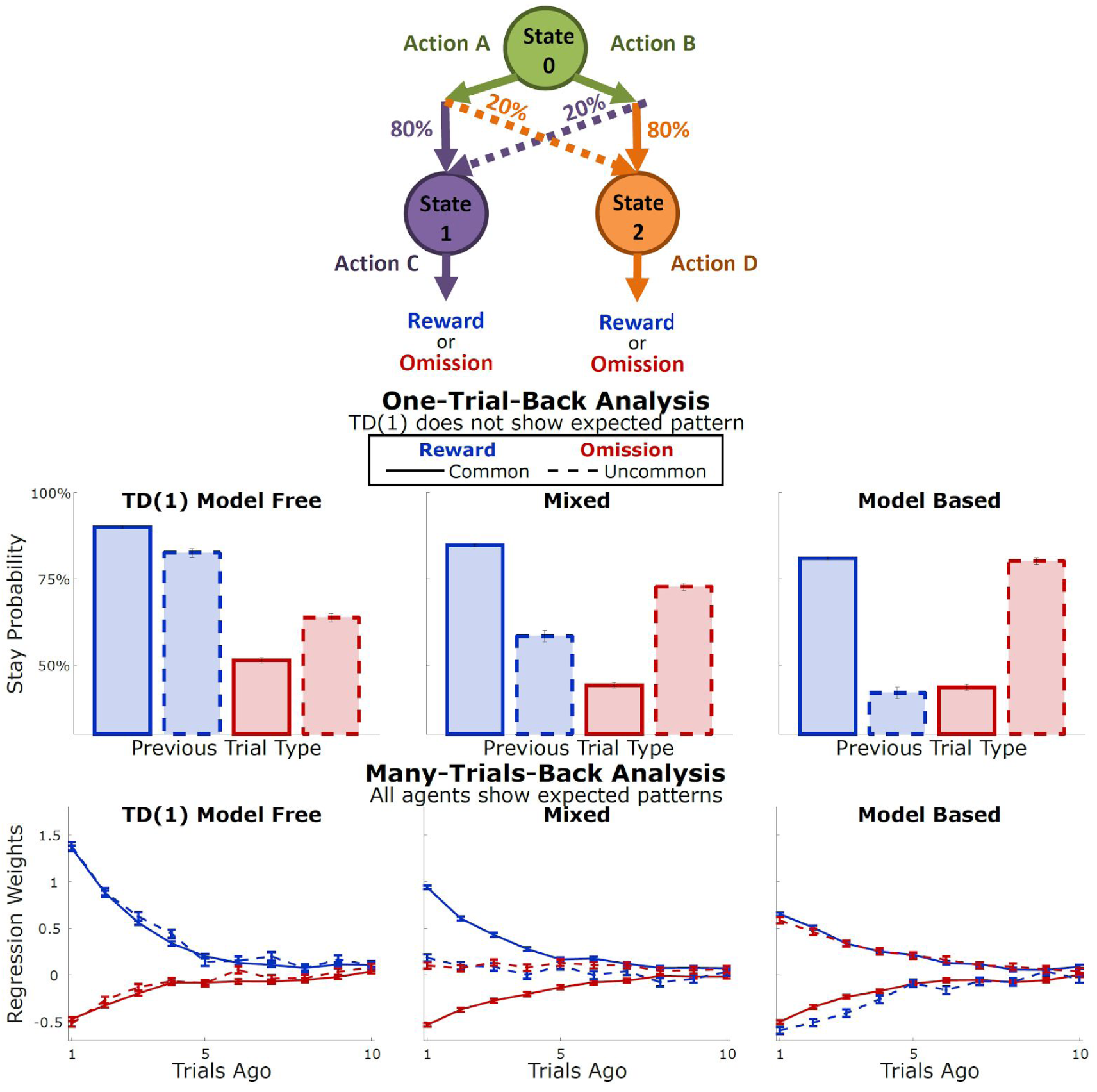
Model-free and model-based patterns of behavior on an alternate version of the two-step task optimized for use with animals.

### Synthetic Behavioral Datasets

To test our analytical techniques, we generated a variety of synthetic datasets for which the “ground truth” generative mechanism is known. In Part I, we analyze synthetic datasets from model-based, model-free, and hybrid agents performing both versions of the task described above (for an introduction to reinforcement learning agents, see Sutton & Barto, 1998). Our model-free agents implement a SARSA(1) algorithm which assigns values to each first-step choice based on the frequency with which it has led to a reward vs. an omission in the recent past. Our model-based agents are aware of the transition probabilities in the task, and use this knowledge to assign values to the choices in accordance with task structure. Hybrid agents mix both of these strategies according to a weighting parameter *w*. These agents are all special cases of the model used in Daw et al. (2011) being run in generative mode: our model-free agents have the parameter *w* set to 0, our model-based agents have *w* set to 1, and all agents in Part One have the parameter *λ* set to 1.

In Part Two, we analyze datasets from agents using three types of alternate strategies. The first type of agents are model-free SARSA(*λ*) agents, incorporating an additional model-free learning rule at the time of the second step transition, in addition to the SARSA(1) learning rule at the time of the final outcome. These are equivalent to the model used in Daw et al., (2011) with the parameter *w* set to 0, allowing the parameter *λ* to vary. The second type are model-free SARSA(1) agents operating on an expanded state space (Akam, Costa, and Dayan 2015) – instead of considering each trial to begin in the same state, these agents consider the initial state of a trial to be different depending on the outcome of the previous trial. The third type are model-based agents incorporating a learning rule on the model itself. These agents maintain a running estimate of how likely it is that each second-step state will follow each first-step action, rather than taking advantage of a static model.

### One-Trial-Back Stay/Switch Analysis

The one-trial-back stay-switch analysis is the most widely used method for characterizing behavior on the two-step task. This method quantifies the tendency of an agent to repeat the choice that it made on the last trial vs. switch to the other choice, as a function of the last trial’s outcome. We consider four possible outcomes: reward-common (RC), reward-uncommon (RU), omission-common (OC), and omission-uncommon (OU). We plot the fraction of trials in each category following which the agent repeated its choice, along with a 95% confidence interval. One-trial-back model-based and model-free indices are computed from the stay probabilities following each outcome as follows:

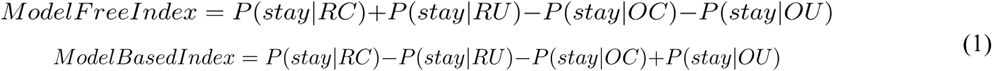

### Many-Trials-Back Regression Analysis

We quantify the effect of past trials and their outcomes on future decisions using a logistic regression analysis based on previous trials and their outcomes (Lau and Glimcher 2005). We define vectors for each of the four possible trial outcomes (RC, RU, OC, OU as above), each taking on a value of+1 for trials of their type where the rat selected the left choice port, a value of −1 for trials of their type where the rat selected the right choice port, and a value of 0 for trials of other types. We define the following regression model:

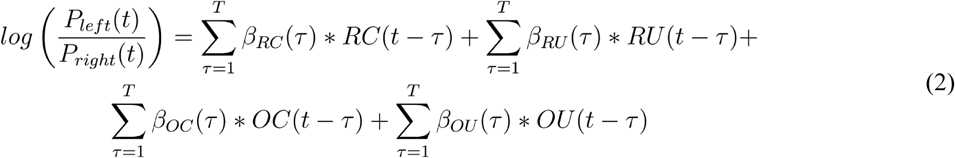

where *β_RC_*, *β_OC_*, *β_RU_*, and *β_OU_* are vectors of regression weights which quantify the tendency to repeat on the next trial a choice that was made *τ* trials ago and resulted in the outcome of their type, and *T* is a hyperparameter governing the number of past trials used by the model to predict upcoming choice. We define a model-based and a model-free index based on sums of fit regression weights consistent with each pattern. This extends the logic of the standard indices (equation 1) to consider many past trials:

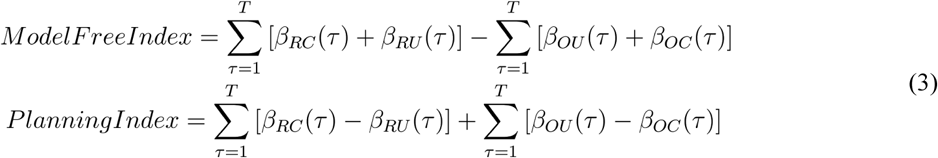

### Model-fitting Analysis

We performed model-fitting analyses in the standard task using the model introduced by Daw et al., (2011). This model has seven parameters, *β*_1_, *β*_2_, *α*_1_, *α*_2_, *λ*,*w*, and *p*. which we allowed to vary, with *α*_1_, *α*_2_, *λ*, and *w* constrained between zero and one. In the simplified task, the parameter *β*_2_ is without meaning and was dropped from the model. In both cases, we performed maximum a posteriori fits using the probabilistic programming language Stan through its MATLAB interface (Carpenter, et al. 2016; Stan Development Team 2016), incorporating weakly informative priors on all parameters (Gelman et al, 2013; *α*_1_ *α*_2_, *λ*, and *w*: beta distribution with a=3, b=3; *β*_1_, *β*_2_, and *p*: normal distribution with μ = 0, σ = 10). Incorporating a prior on the learning rates was necessary in order to avoid pathological fits in which β_1_ and λ become unconstrained as α_1_ goes to zero, or β_2_ becomes unconstrained as α_2_ goes to zero. To compute model-based and model-free indices from these fit parameters, we take advantage of the fact that the parameter *w* controls the relative strength of each pattern, with *w* = 1 being fully model-based and *w* = 0 being full model-free, while *β_1_* controls the overall degree of decision noise on the first step, with lower values corresponding to behavior that is more random. We therefore compute model-based and model-free indices from the values of the fit parameters as follows:

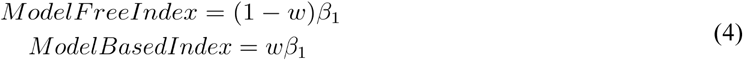

## Results

### Section I: Characterizing Model-Based vs. TD(1) Model-Free Behavior

The two-step task was developed as an experimental tool for separately quantifying model-based and model-free patterns in behavioral data in a trial-by-trial setup. In the first step of the task, the subject is presented with and first-step choice state (state 0 in figure 1), and selects between two available actions (Actions A and B in figure 1). The subject then experiences a probabilistic transition to one of two possible second-step choice states (States 1 & 2). Each first-step action leads with high probability to one of the second-step states and with low probability to the other second-step state. In each second-step state, the subject is presented with a choice between two second-step actions (C & D or E & F). Each of the four second-step actions leads to a reward with a certain probability. These reward probabilities are different from one another, and are changing with time, so optimal performance on the task requires continuous learning about the second-step reward probabilities, and selection of the first-step action which is most likely to lead to the best second-step action becoming available. Optimal behavior therefore requires knowing the relationship between first-step actions and their likely outcomes – that is, optimal behavior on this task requires use of a model.

While optimal performance on the two-step task requires use of a model, a variety of model-free strategies exist as well. The most commonly considered model-free strategy, TD(1), simply learns to repeat first-step actions which are followed by rewards, and to avoid those which are followed by omissions, ignoring the structure of the task. These strategies differ dramatically from model-based strategies in how they alter their choice behavior following uncommon transition trials. In particular, model-free strategies tend to increase their likelihood of repeating a first-step action which is followed by a reward, and to decrease it when it is followed by an omission, regardless of whether the transition was common or uncommon. Model-based strategies, on the other hand, tend to decrease their likelihood of selecting a first-step action that led to a reward after an uncommon transition (because the other first-step action is most likely to lead to the state that preceded that reward), and similarly to increase their likelihood of selecting a first-step action that led to an omission after an uncommon transition (because the other first-step action is most likely to lead to the state that preceded the omission). This difference suggests that analysis of choice behavior should be sufficient to identify a behavioral dataset as arising from either a model-based or a model-free strategy.

The simplest and most common way of analyzing choice behavior is a one-trial-back analysis of stay/switch behavior. In this analysis, the dataset is divided into four sets of trials based on first-step transition (common/uncommon) and second-step outcome (reward/omission), and analyzed as to whether the first-step choice on the following trial was the same (stay) or different (switch). The prediction is that model-free agents will show larger stay probability following rewards and lower stay probability following omissions, regardless of transition, while model-based agents will show larger stay probability following reward-common and omission-uncommon trials, and lower stay probability following omission-common and reward-uncommon. This prediction is borne out in synthetic datasets generated by TD(1), model-based, and hybrid agents (Fig 1, middle). A common way of summarizing these data is to compute a “model-free index” as the main effect of reward (reward/omission) on stay probability, and a “model-based index” as the reward by transition (reward/omission by common/uncommon) interaction effect (see Eq. 1).

To provide a richer picture of patterns in behavior, we consider a “many-trials-back” analysis which takes into account multiple past trials. This analysis is a logistic regression, including separate weights for each possible outcome (common/uncommon, reward/omission) at each trial lag (see Method, eq. 2). In these models, we expect the same patterns that we expected from the one-trial-back analysis: model-free systems will tend to increase their probability of repeating a choice after rewards and decrease it after omissions, independently of transition, while model-based systems will show opposite patterns after common vs. uncommon trials. We see that the many-trials-back analysis applied to synthetic data from TD(1), model-based, and hybrid agents indeed produce these patterns (Fig 1, below).

In addition to revealing more information about patterns in behavior, the many-trials-back analysis has the advantage of robustness to changes in task design. In figure 2, we consider an alternative version of the two-step task optimized for use with rodents. In this version of the task, the second-step choice is eliminated, and each second-step state contains only one available action. The transition probabilities and reward probability dynamics are also slightly different (see *Method, Two-Step Behavioral Task* for details). The one-trial-back analysis applied to synthetic data from this task reveals a strong difference in stay probability following reward-common vs. reward-uncommon trials, as well as following omission-common vs. omission-uncommon for a model-free TD(1) agent (Figure 2, middle row, left). This analysis would result in a positive model-based index for this purely model-free agent, an undesirable result. This happens because the one-trial-back analysis considers only the outcome of the immediately previous trial, while the behavior of the agents is determined in part by outcomes that occurred on trials further in the past (e.g. for an agent with a learning rate of 0.3, the immediately previous trial’s outcome contributes only 30% to the overall value). If the decision of the agent depends on the outcomes of many previous trials, and if the outcome of the immediately previous trial covaries with the outcomes of many previous trials, we expect a one-trial-back analysis to be subject to confounds. The many-trials-back analysis is robust to these issues, since it accounts explicitly for the impact of past trial on decision behavior (Figure 2, bottom row).

These effects are most acute for low learning rate agents, since these agents’ behavior is most dominated by events more than one trial in the past. To illustrate this systematically, we generated synthetic datasets at a variety of learning rates, and computed the model-free and model-based index for each using the one-trial-back method (Figure 3, top row). For the model-free agent, we found that the model-free index increased substantially with learning rate, while the model-based index decreased slightly. For the model-based agent, the model-based index increased substantially with learning rate, while the model-free index was relatively unaffected. For the mixed agent, both indices increased with learning rate. We find that model-based and model-free indices computed using the many-trials-back method (Figure 3, middle row), are affected only slightly by changes in learning rate. We also generated synthetic datasets from mixed agents in which we systematically varied the learning rate of either the model-based agent (Figure 4, top), or the model-free agent (Figure 4, bottom). We find that indices computed by the one-trial-back method are separately affected by these changes -- this can result in datasets generated by mechanisms that are equal mixtures of model-based and model-free systems being characterized as either predominantly model-free or predominantly model-based, depending on learning rate (Figure 4 left, orange and green lines cross as learning rate changes). Indices computed by the many-trials-back method are much less sensitive to changes in one agent’s learning rate (Figure 4 second from left).

**Figure 3:**
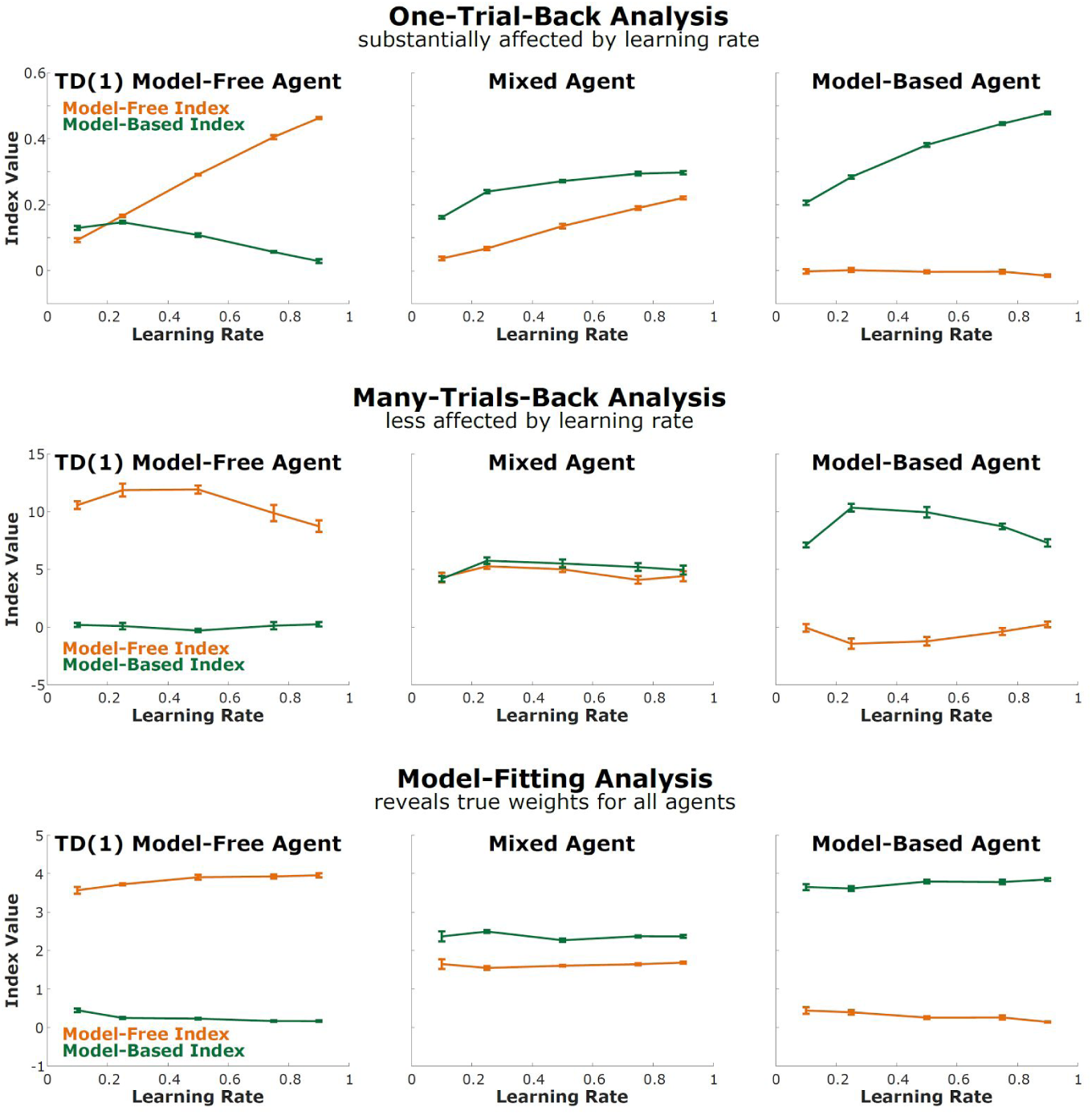
Changing the learning rate. Model-based and model-free indices produced by the one-trial-back (above), many-trials-back (middle) and model-fitting method (below), applied to datasets from TD(1) model-free agents, model-based agents, and mixed agents with different learning rates.

Another way of common way of characterizing behavior on the two-step task involves fitting the parameters of a particular agent-based computational model (Daw, et al., 2011) to the behavioral dataset. One of these parameters, *w*, controls the extent to which a dataset is dominated by model-based vs. model-free patterns. We performed maximum a posteriori fits of this model to our synthetic datasets, and recovered parameter estimates. While it is typical in the literature on this task to simply interpret the weighting parameter w as a relative measure of the strength of model-based vs. model-free patterns, here we combine *w* and *β*_1_ to compute separate model-based and model-free indices (see *Method*, eq. 4), for simpler comparison to the one-trial-back and many-trials-back methods. One assumption of this model is that the model-based and model-free agents have the same learning rate as one another. It therefore does a good job of characterizing the behavior of agents for which this assumption is met, reporting model-based and model-free indices that are robust to changes in learning rate that affect both systems equally (Figure 3, bottom). It does much less well at characterizing agents for which this assumption is not met (Figure 4, second from right; note that orange and green lines cross). If we generalize the model to relax this assumption, it gains the ability to characterize these agents correctly (Figure 4, right). The fit parameters of agent-based models, therefore, can be a powerful tool for understanding the mechanisms that gave rise to a behavioral dataset, performing better than the other tools we consider here (Figure 3, bottom; figure 4, right). In cases where the model makes assumptions that are not met, however, these fits can be misleading (Figure 4, second from right).

**Figure 4:**
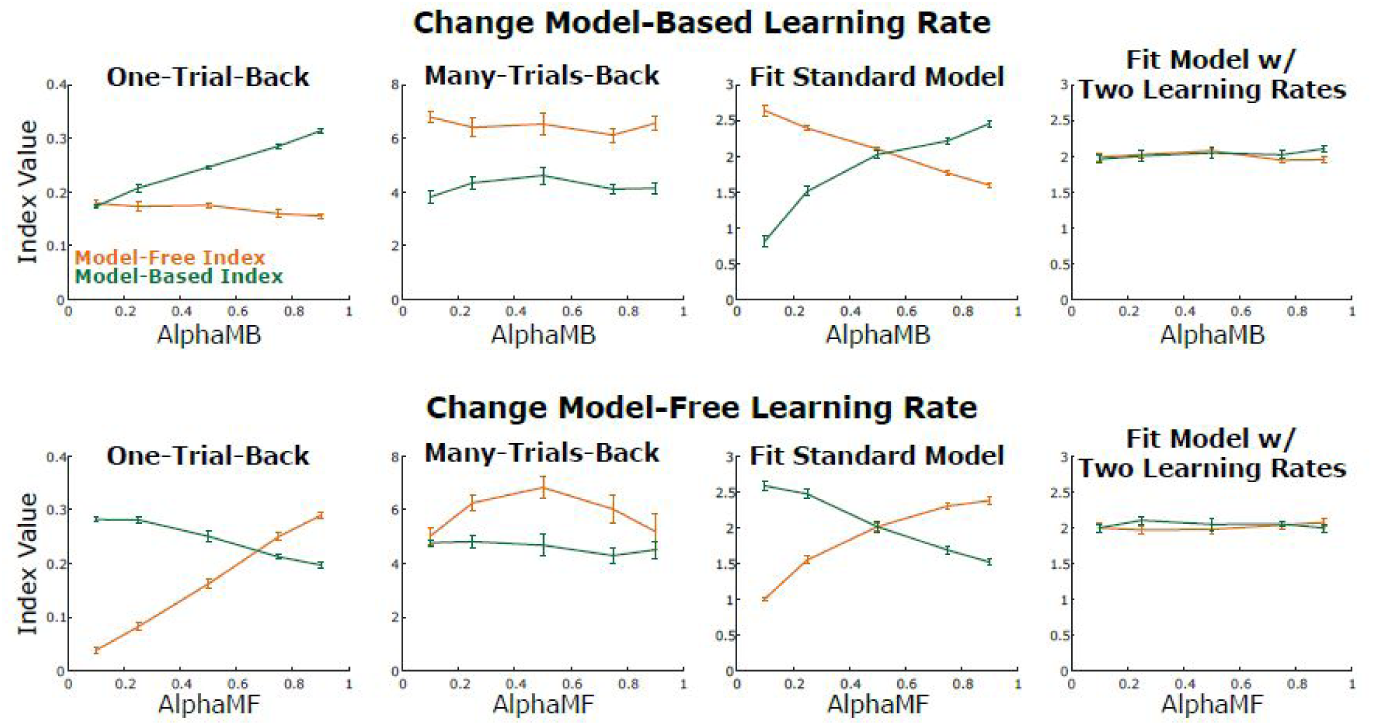
Changing one learning rate at a time. Model-based and model-free indices produced by the one-trial-back, many-trials-back, standard model fit, and expanded model fit.

In this section, we have considered the case of synthetic behavioral datasets generated by a particular model-based algorithm, a particular model-free algorithm, and hybrid systems mixing both together. With respect to these datasets, we have shown that the one-trial-back analysis is a relatively unreliable tool for characterizing behavioral patterns, but that both the many-trials-back analysis and explicit fits of the generative model are more useful. In cases where the assumptions built into a model are true of the process which generated a dataset, fits of that model are ideal for characterizing the dataset (Figure 3, bottom; figure 4, right). In cases where these assumptions are not met, model fits can be badly misleading (Figure 4, second from right). In the next section, we consider synthetic behavioral datasets generated by different types of algorithms. We will show that only fits of agent-based models are able to properly characterize these datasets in general, but that the assumptions built into these models can to some extent be checked using tools like the many-trials-back regression analysis.

### Section II: Strategies beyond simple MB/MF

In Section I, we showed that both the many-trials-back regression analysis as well as fits of agent-based computational models are effective techniques for characterizing particular kinds of behavioral datasets from the two-step task. In this section, we will show that the many-trials-back analysis is able to provide a relatively rich and theory-neutral description of patterns present in a behavioral dataset, at the expense of providing relatively little insight into the mechanisms which may have generated those patterns. Agent-based models, on the other hand, provide theoretical insight at the expense of relying on theoretical assumptions – if these assumptions are not met, the description of behavior provided by a model fitting approach will be at best incomplete, and at worst actively misleading. That theory-neutral approaches like the many-trials-back analysis can reveal when the assumptions of a model are violated, providing important guidance to a model fitting analysis approach.

First, we consider a class of model-free systems known as TD(λ) systems, which are widely used in psychology and neuroscience as models of human and animal behavior. In addition to reinforcing first-step actions when they lead to reward, these systems introduce an extra learning rule reinforcing first-step actions when they lead to second-step states which themselves have led to reward on previous trials. These algorithms introduce an “eligibility trace” parameter λ to control the relative importance of these two updates. This λ-modulated double update rule is incorporated into the standard model used in the literature to characterize behavioral data on this task. We generated synthetic datasets from systems with different values of λ, and subjected these datasets to the many-trials-back analyses (Figure 5, above). We see that as λ decreases, a characteristic pattern arises which is qualitatively different from that pattern shown by any of the agents considered earlier. This pattern involves reward by transition interactions at long trial lag, giving rise to erroneous “model-based indices” > 0 for these datasets analyzed using the many-trials-back method. We compute model-based and model-free indices using the one-trial-back, many-trials-back, and model-fitting methods, and see that only agent-based model fitting is robust to changes in lambda.

**Figure 5:**
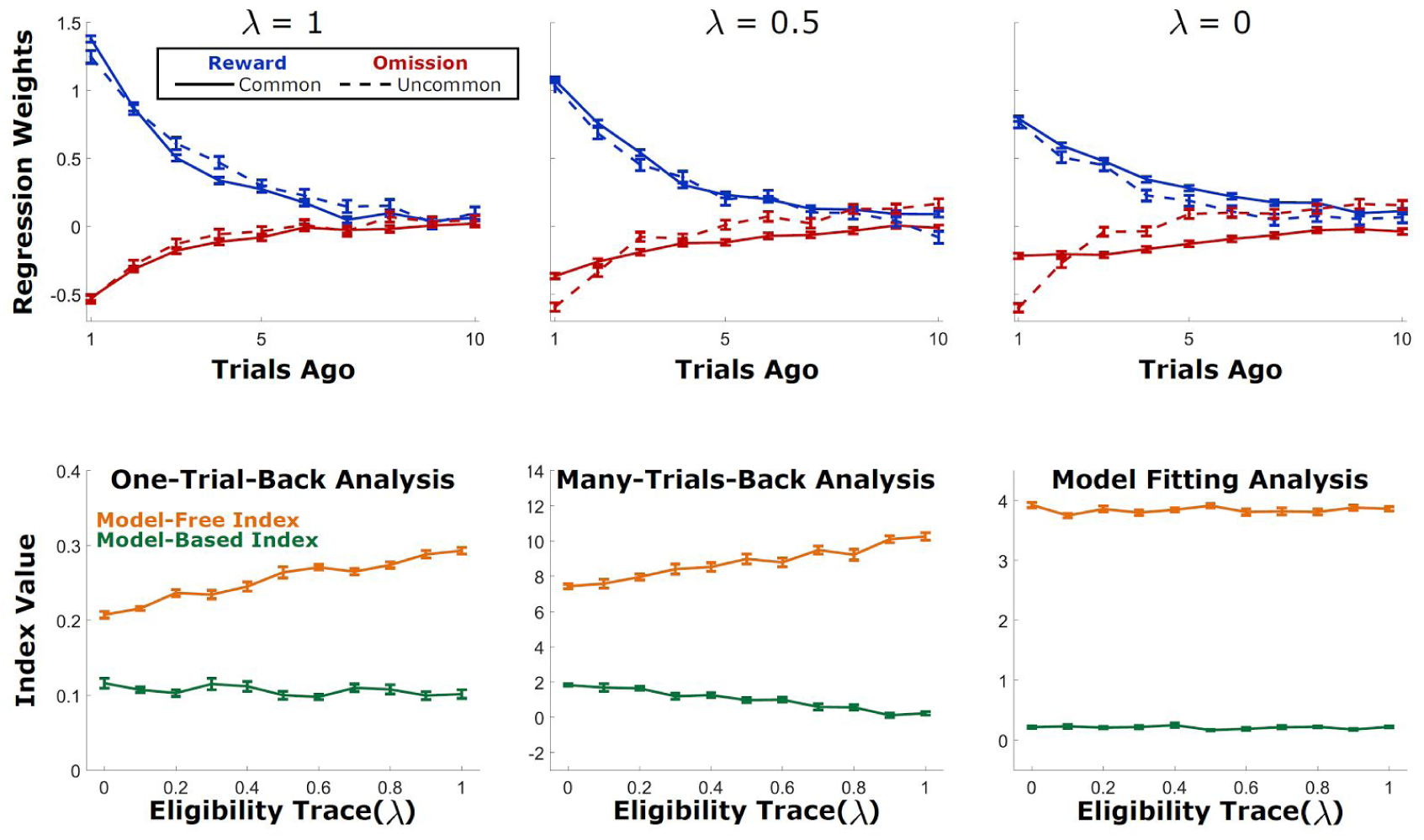
Changes in eligibility trace parameter confound both the one-trial-back and the many-trials-back analyses.

Explicit model-fitting is robust to changes in both alpha and lambda, and is a robust and statistically efficient way to characterize behavior that was generated by a process which meets the theoretical assumptions built into the model. Model fitting can lead to confusing or misleading results if applied to a dataset which does not meet these assumptions. Next, we consider two such types of systems. The first are model-free systems which take advantage of an expanded state space (Akam, Costa, and Dayan 2015). We term these system “expanded state space model-free” or ESS-MF. Instead of considering each trial to begin in the same state (“State 0” in figure 1), they consider different trials to begin in different states, according to the outcome of the previous trial. They therefore learn different values of actions A&B for each of these possible states. The “state space” considered by one of these agents is determined by a particular set of features of the previous trial that it takes into account. In figure 6, we consider an agent taking into account the previous trial’s outcome and reward (ESS-MF OR, four total states), as well as one considering the previous trial’s choice, outcome, and reward (ESS-MF COR, eight total states), and show the results of the one-trial-back, many-trials-back, and model-fitting analysis to a synthetic dataset. This dataset shows large reward by transition interaction effects (“model-based pattern”) in the one-trial-back analysis and in the many-trials-back analysis at lag one, but reveals main effects of reward (“model-free pattern”) at longer trial lags.

**Figure 6:**
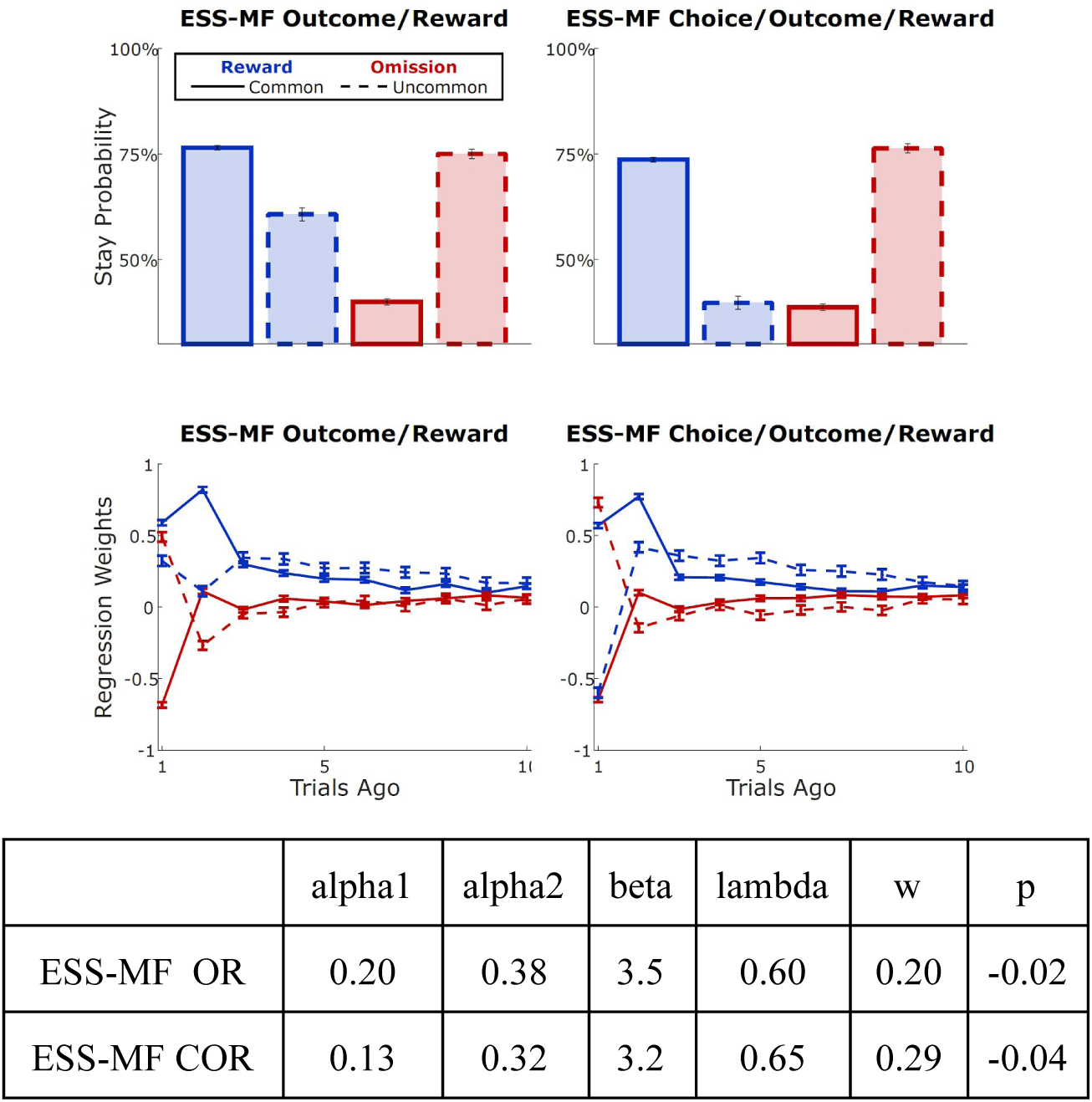
“Enhanced state-space” model-free systems mimic model-based or hybrid systems in the one-trial-back analysis, reveal distinctive patterns in the many-trials-back analysis.

Fitting the standard model to these datasets results estimates for the *w* parameter that are greater than zero (w=0.20 and 0.29 for the outcome-reward and choice-outcome-reward versions, respectively), erroneously suggesting that the dataset was generated in part by a model-based mechanism. The model-fitting analysis on its own reveals no insight into the fact that its assumptions are not met by these particular datasets: Indeed, it is blind to the difference between these datasets and ones generated by a hybrid MB/TD(λ) system using the estimated parameters. We illustrate this by generating such datasets and performing the model-fitting and many-trials-back analyses. While the model-fitting recovers very similar parameter estimates, the many-trials-back analysis reveals qualitative differences between the datasets (compare figure 6 middle and bottom rows to figure 7), pointing to the fact that the original dataset was generated by mechanisms that violate the assumptions of the fitting analysis.

**Figure 7:**
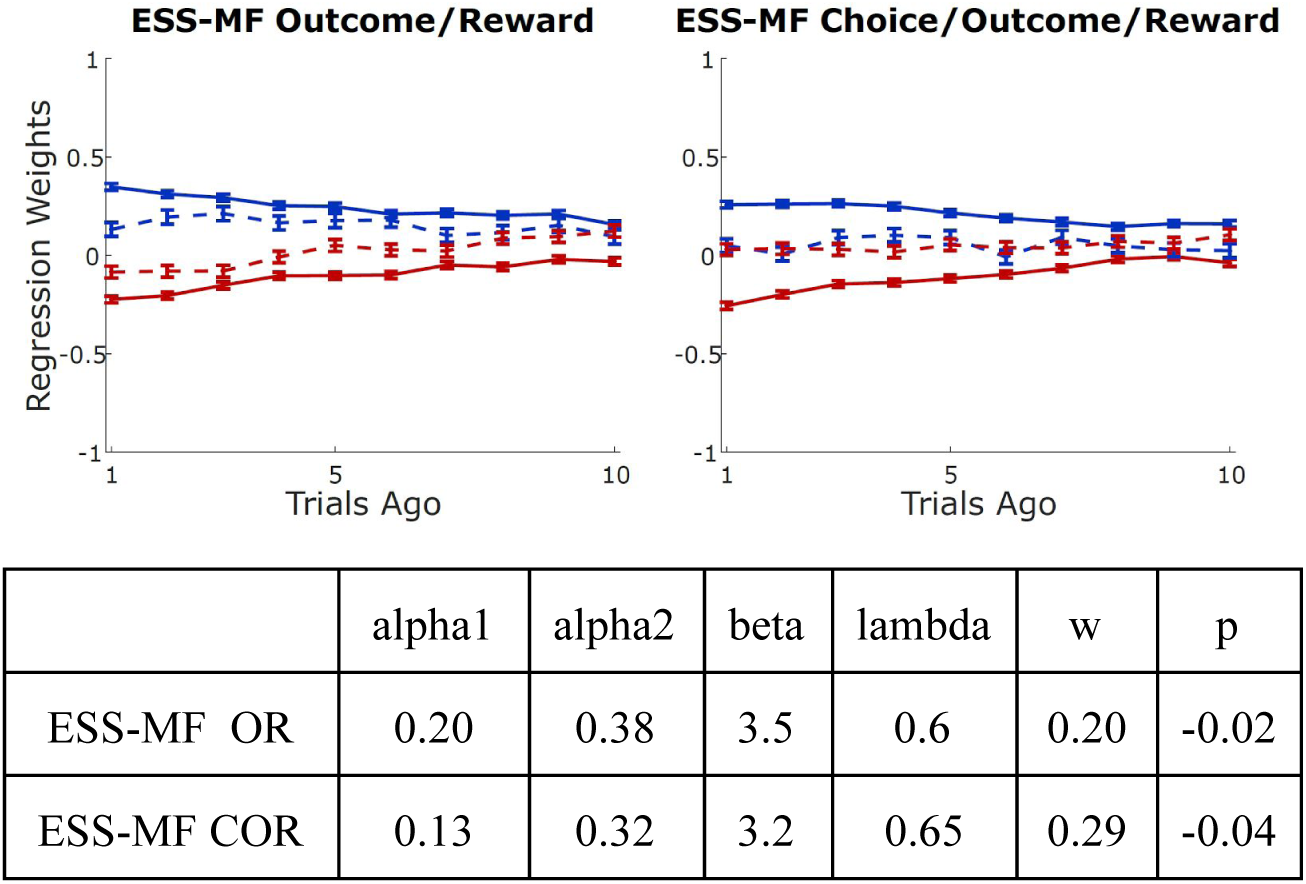
Datasets generated using the standard model with parameters fit to ESS-MF datasets. Many-trials-back analysis reveals qualitative differences between these datasets and the EFF-MF datasets, revealing that the assumptions of the standard model do not apply to this dataset. Fits of the standard model itself reveal no differences.

Finally, we consider model-based systems which also violate the assumptions of the fitting analysis. Instead of having access to a perfect and stable model of the world, these systems learn their model from experience and update the model as new information becomes available. Figure 8 (top) shows the results of the many-trials-back analysis applied to a synthetic dataset from two agents with different model-learning rates. This analysis reveals differences between the effects of common and uncommon trials (solid and dotted lines), which would result in a nonzero “model-free index” for this purely model-based system. We illustrate this systematically, changing the learning rate with which the agent updates its model of the world from 0 to 0.5, and computing model-based and model-free indices using the one-trial-back, many-trials-back methods, as well as fits of the standard agent (which does not update its model) and a generalized agent which does update its model. Only the generalized agent reports indices which are accurate: a model-free index near zero and a model-based index that is stable with respect to changes in learning rate.

**Figure 8:**
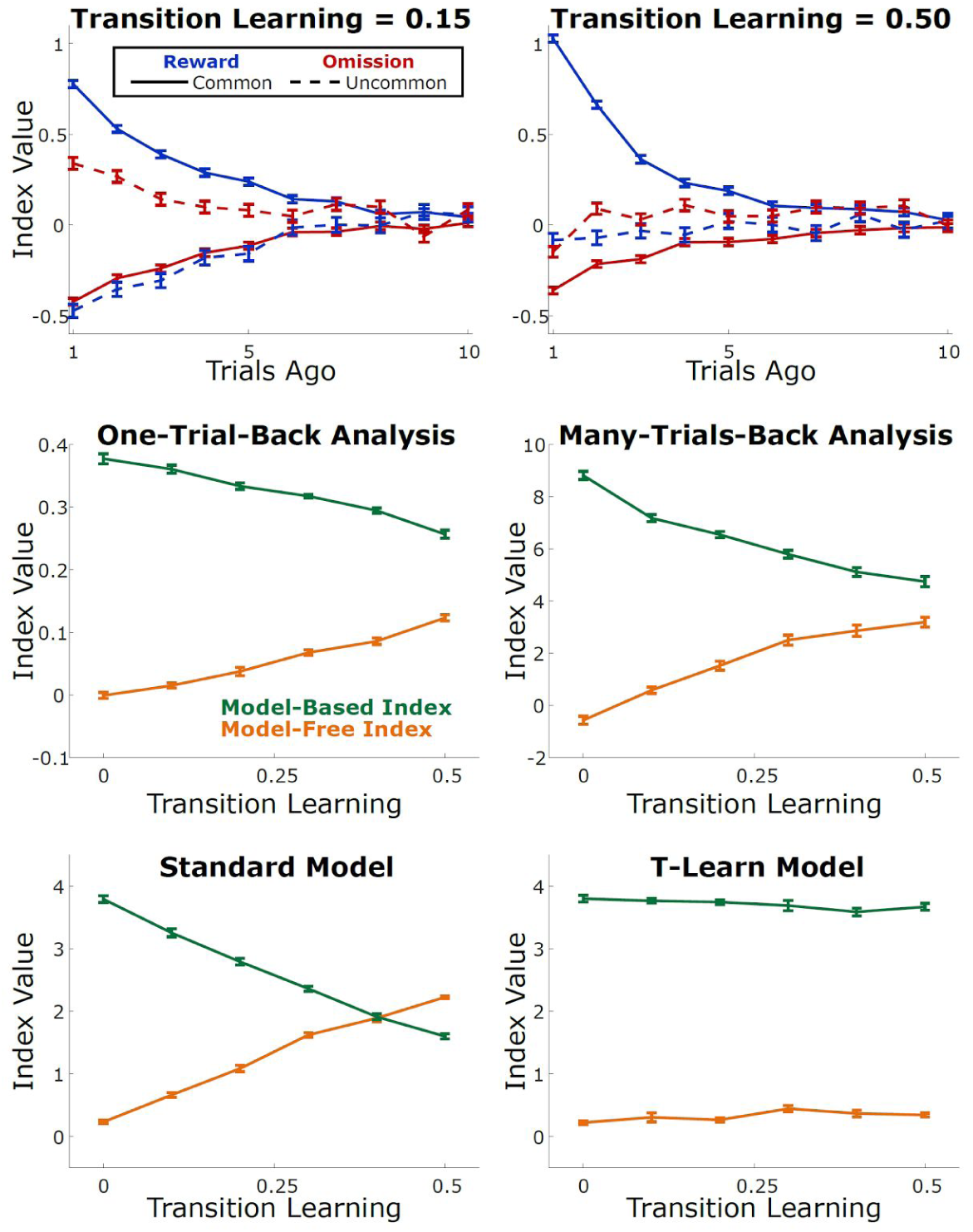
Datasets generated by an agent with a learning rule for the transition probabilities. Many-trials-back analysis reveals qualitative patterns that are difference from other agents we have considered (top row). Indices computed using one-trial-back, many-trials-back, or standard model are misleading, while indices computed by fitting a model that has been generalized to consider transition learning are not.

## Conclusions

The recently developed “two-step task” of Daw et al., (2011) has become a popular tool for investigating the psychological and neural mechanisms of planning, promising a behavioral readout of planning in the context of a repeated trials decision making task. Crucial to its success is the idea that analysis of behavior can reveal the extent to which a subject is performing the task using a planning vs. a model-free strategy. Work on this task typically uses two tools for characterizing these patterns: a relatively theory-neutral one-trial-back analysis of choice behavior, and parameter estimation using a particular computational model, both of them introduced in the pioneering work on this task by Daw et al (2011). In the present work, we investigated the performance of each of these tools, and compared them to a novel many-trials-back method. We showed that in a modified version of the task, the one-trial-back analysis fails to adequately characterize behavior even for relatively simple agents, with particularly acute difficulties categorizing model-free systems with low learning rates (Fig 3). This could lead to a behavioral dataset produced by a low learning rate model-free agent being misclassified as having come from a hybrid system. The absolute results of the one-trial-back analysis vary strongly with learning rate for agents of all types (Fig 3), a result with implications for studies investigating group or individual differences in the balance between model-based and model-free control. The model-fitting method is suitable for characterizing patterns in behavioral datasets which were generated by mechanisms conforming to its assumptions (Fig 3 & 4), but provides at best an incomplete account of behavior in cases where those assumptions were not met (Figs 5 & 7).

We introduce a novel many-trials-back method, which alleviates some of the problems with the one-trial-back method, and provides a valuable complement to agent-based model fitting. This method is able to accurately categorize simple hybrid agents in a manner relatively insensitive to learning rate (Fig 3). It provides a rich description of the patterns present in behavior, and can be used to discriminate datasets generated by simple hybrid agents from those generated by alternative mechanisms including SARSA(λ), enhanced state space SARSA(1), and MB with model update. This puts the many-trials-back analysis in a position to be useful for model checking (Gelman et al. 2013).

In complex tasks like the two-step task, many behavioral strategies are possible. In general, it is challenging to devise theory-neutral analyses which distinguish these strategies, since very different strategies can give rise to similar qualitative patterns of behavior. Fits of agent-based models can rise to this challenge, distinguishing strategies based on quantitative differences between the patterns of behavior they produce. These models, however, rely on assumptions about the strategy that gave rise to a dataset, and if these assumptions are not met, their fit parameters can be difficult to interpret. One way to check whether the assumptions that underlie a particular agent-based model are true about a particular dataset is to fit the model to a dataset, generate a synthetic dataset using the fit parameters, and compare this synthetic dataset to the actual dataset using rich and theory-neutral analysis methods.

We recommend that future work on the two-step task use caution in interpreting the results of the one-trial-back analysis, and where possible take advantage of approaches like the many-trials-back analysis, both to characterize behavior in a theory-neutral way, and to check the assumptions built into analysis using agent-based models.

